# High nucleotide substitution rates associated with retrotransposon proliferation drive dynamic secretome evolution in smut pathogens

**DOI:** 10.1101/2021.04.23.441129

**Authors:** JRL Depotter, B Ökmen, MK Ebert, J Beckers, Kruse Jb, M Thines, G Doehlemann

**Author notes:** Corresponding author (GD).

## Abstract

Transposable elements (TEs) play a pivotal role in shaping diversity in eukaryotic genomes. The covered smut pathogen on barley, *Ustilago hordei*, encountered a recent genome expansion. Using long reads, we assembled genomes of 6 *U. hordei* strains and 3 sister species, to study this genome expansion. We found that larger genome sizes can mainly be attributed to a higher genome fraction of long terminal repeat retrotransposons (LTR-RTs). In the studied smut genomes, LTR-RTs fractions are the largest in *U. hordei* and are positively correlated to the mating-type locus sizes, which is up to ∼560 kb in *U. hordei*. Furthermore, LTR-RTs were found to be associated with higher nucleotide substitution levels, as these higher levels occur more clustered in smut species with a recent LTR-RT proliferation. Moreover, genes in genome regions with higher nucleotide substitution levels generally reside closer to LTR-RTs than other genome regions. Genome regions with many nucleotide substitutions encountered an especially high fraction of CG substitutions, which is not observed for LTR-RT sequences. The high nucleotide substitution levels particularly accelerate the evolution of secretome genes, as their more flexible nature results that substitutions often lead to amino acid alterations.

**Importance:** Genomic alteration can be generated through various means, in which transposable elements (TEs) can play a pivotal role. Their mobility causes mutagenesis in itself and can disrupt the function of the sequences they insert into. Indirectly, they also impact genome evolution as their repetitive nature facilitates non-homologous recombination. Furthermore, TEs have been linked to specific epigenetic genome organizations. We report a recent TE proliferation in the genome of the barley covered smut fungus, *Ustilago hordei.* This proliferation is associated with a distinct nucleotide substitution regime that has a higher rate and a higher fraction of CG substitutions. This different regime shapes the evolution of genes in subjected genome regions. Our findings highlight that TEs may influence the error-rate of DNA polymerase in a hitherto unknown fashion.

## INTRODUCTION

Transposable elements (TEs) play a pivotal role in the genome evolution of eukaryotic organisms, including fungi (1). Fungal genomes can vary considerably in size, which is often determined by the extent and recency of TE proliferations (2, 3). On one side of the spectrum, Microsporidia, a diverse group of obligate intracellular parasitic fungi, contain members with extremely small genomes down to 2.3Mb that lack TEs (4, 5). In contrast, rust plant pathogens from the order Pucciniales contain members with genome sizes that are among the largest in the fungal kingdom (6, 7). For instance, the wheat stripe rust pathogen *Puccinia striiformis* f.sp. *tritici* has an estimated genome size of 135 Mb, which more than half consists of TE sequences (8). Mutations caused by TE transposition predominantly have a neutral or negative impact, but in particular cases they can also improve fungal fitness (3, 9). For plant pathogenic fungi, TE transposition can be a source of mutagenesis to evade host immunity and/or lead to an optimized host interaction (10). TEs can also passively contribute to mutagenesis, as their transpositions increase homologous genome sequences that are prone to ectopic recombination (11, 12). Pathogens evolve by host jumps, radiation and subsequent arms races with their hosts (13), in which the latter attempts to detect pathogen ingress through the recognition of so-called invasion patterns (14). One invasion pattern that is typically detected are effectors, i.e. secreted proteins that facilitate host colonization (15). To quickly adapt to effector-triggered immunity and yet continue host symbiosis, effector genes often reside in genome regions that facilitate mutagenesis (13, 16), such as those rich in TEs (17). TE-rich genome regions may not only encounter higher mutation rates, but may also have a higher chance to fix mutations due to their functionally more accessory nature (18).

TEs are a diverse group of mobile nucleotide sequences that are categorized into two classes (1). Class I comprises retrotransposons that transpose through the reverse transcription of their messenger RNA (mRNA). Class II are DNA transposons that transpose without mRNA intermediate. TEs are then further classified based on their sequence structure (19). Retrotransposons with direct repeats at each end of their sequence are long terminal repeat retrotransposons (LTR-RTs) (20). LTR-RTs can encode the structural and enzymatic machinery for autonomous transposition. However, they may lose this ability through mutagenesis, but still be able to transpose using proteins of other TEs (21). LTR-RTs can then be further classified into superfamilies including *Copia* and *Gypsy*, which differ in the order of their reverse transcriptase and their integrase domain (19).

Smut fungi are a diverse group of plant pathogenic, hemibiotrophic basidiomycetes of which many infect monocot plants, in particular grasses. They live saprophytically as yeasts and mate in order to switch to the diploid, filamentous stage that enables them to infect their host (22). Smut pathogens are very host-specific and generally have small genomes in comparison to other plant pathogens (23, 24). The corn smut species *Ustilago maydis* and *Sporisorium reilianum* are closely related and have genome sizes of 19.8 Mb and 18.4 Mb, respectively (25, 26). This is partly due to their low level of repetitive sequences, including TEs. In total, only 2.1 and 0.5% of the genome consists of TEs for the *U. maydis* and *S. reilianum*, respectively (27). The covered smut pathogen of barley, *Ustilago hordei*, and the Brachipodieae grass smut, *U. brachipodii-distachyi*, are two related smut fungi and have genome assemblies of 21.15 Mb and 20.5 Mb, respectively (27, 28). These larger genome assembly sizes correlate to their higher TE content, which is 11.8% and 14.3% for *U. hordei* and *U. brachipodii-distachyi*, respectively (27). The assembled genome of *U. brachipodii- distachyi* is originally published under the species name *U. bromivora* (27). *U. brachipodii- distachyi* infects members from the tribe Brachipodieae, whereas *U. bromivora* affects bromes from the supertribe Triticodae (29, 30). Considering the host specific nature of smut pathogens, we prefer to refer to this assembly as *U. brachipodii-distachyi* instead of *U. bromivora*, as the assembled strain infects *Brachypodium* species (27, 29).

Mating in grass-parasitic smut fungi is tetrapolar in *U. maydis* and *S. reilianum*, whereas *U. hordei* and *U. brachipodii-distachyi* have a bipolar mating system. In the bipolar system, there is one mating-type locus where recombination is suppressed (27, 31, 32). This locus is flanked by the *a* locus, which contains pheromone/receptor genes, and the *b* locus, which encodes the two homeodomain proteins bEast and bWest (33, 34). In the bipolar mating-type system, there are two mating-type alleles, *MAT-1* and *MAT-2*, which are in *U. hordei* ∼500kb and ∼430kb in size, respectively (32). A large fraction of the mating-type loci consists of repetitive sequences, i.e. ∼45% repeats for *U. hordei* (28, 35). In contrast, the tetrapolar smuts, *U. maydis* and *S. reilianum*, have their *a* and *b* loci on different chromosomes causing them to segregate independently during meiosis (26, 31).

Recently, the complete genome of the reference *U. hordei* strain Uh364 was re- sequenced using the long-read PacBio technology and, instead of the previous 21.15 Mb assembly (28), a 27.1 Mb assembly was obtained (36). Thus, the *U. hordei* genome underwent a genome expansion as it is significantly larger than other sequenced smut species (27). This finding triggered us to study the *U. hordei* genome more in depth and use recently developed long-read sequencing technologies to unravel how its genome recently expanded.

## Results

### LTR-RTs is an important determinant for *U. hordei* genome size

To study the expansion in genome size of *U. hordei*, we sequenced and assembled 6 *U. hordei* strains of different geographic origins (Figure S1). Five contained a *MAT-1* locus and one (Uh1278) a *MAT-2.* The assemblies were composed of 23-46 contigs and ranged from 25.8-27.2 Mb in size (Table 1). Strain Uh805 was assembled into 23 contigs that are homologous to the 23 chromosomes of *U. brachipodii-distachyi* indicating that inter-chromosomal rearrangements did not occur during the divergence of these two species (27). *MAT-1* loci, regions between and including the *a* and *b* loci, ranged from 536 to 564 kb in size, whereas the *MAT-2* locus was 472 kb (Table S1). In the *U. hordei* genome assemblies, class I TE sequences are over 6 times more abundant than class II TE sequences (Table 1, S2). More than 90% of the class I TEs consist of LTR-RTs, which is a total sequence amount ranging between 4,326 and 5,272 kb (Table 1). The number of LTR-RT sequences is positively correlated to the assembly sizes (*r* = 0.94, *p*-value = 0.0051). Moreover, using strain Uh805 as a reference, 56-79% of the differences in assembly size with other strains can be attributed to differences in LTR-RT content. Thus, the variation in genome size between *U. hordei* strains can largely be attributed to intraspecific differences in LTR-RT proliferation and/or retention. More than 75% of the mating-type loci consist of repetitive sequences and over 29% are classified as LTR-RTs. The *MAT-1* and *MAT-2* loci and their flanking regions only have 27% one-to-one best homology to each other (Figure 1A), which is mainly due to mating-type specific repeats as only 6% of the repeats are shared between the two mating types. In contrast, 41 of the 47 expressed mating-type locus genes are shared between the two alleles (Figure 1B). Homologous recombination is suppressed in the mating-type region, which makes that TE transpositions within these regions are by definition mating-type specific (Figure 1B) (32).

**Figure 1:**
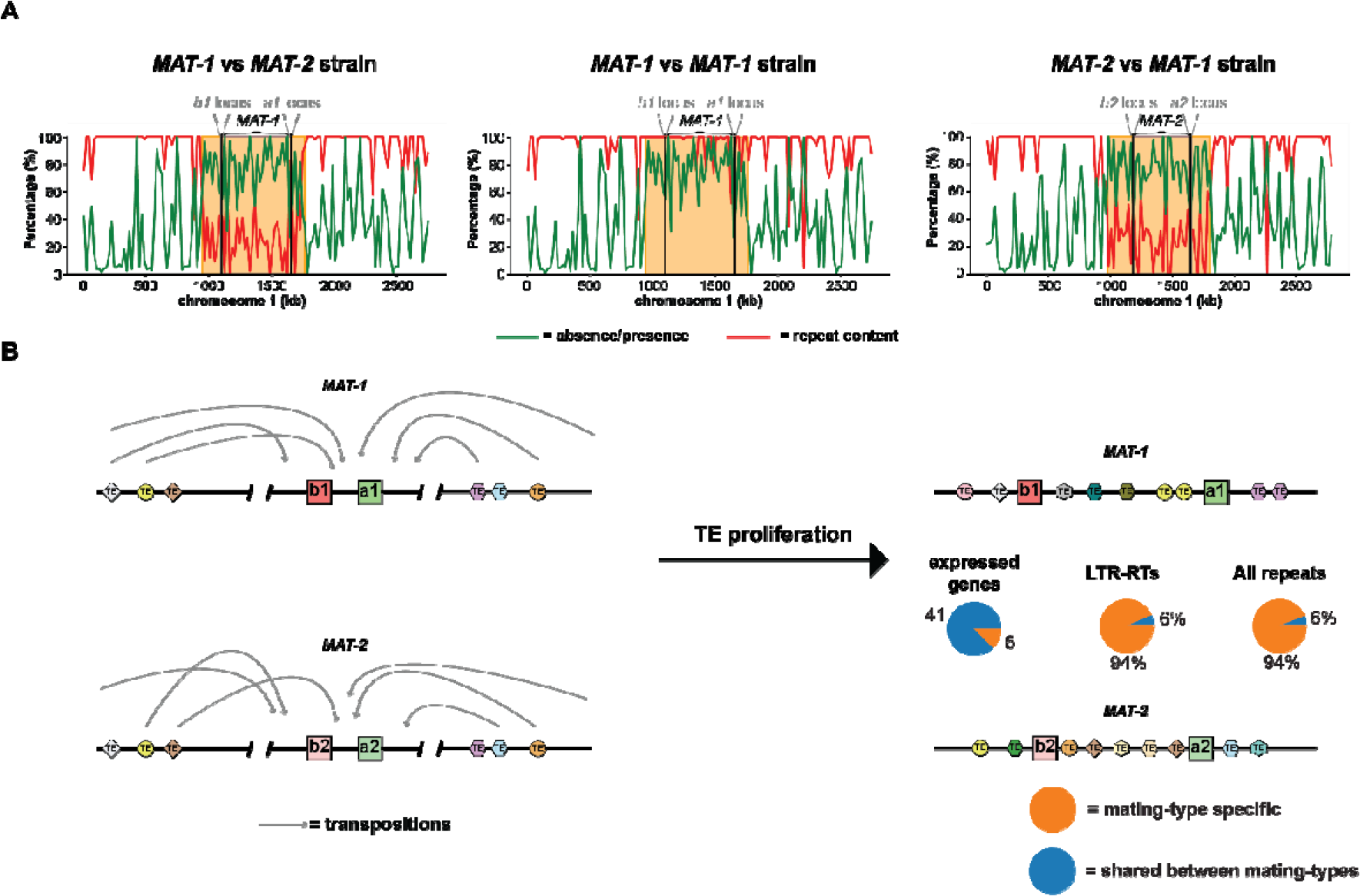
The mating-type specificity of MAT-1 and MAT-2 loci sequences. (A) As references, the MAT-1 strain Uh805 and the MAT-2 strain Uh1278 were used. Repeat content and presence/absence polymorphisms were calculated for 20 kb windows. Presence-absence polymorphisms were determined between the *MAT-1* and *MAT-2* reference strains, in addition to the *MAT-1* strains Uh805 and Uh811. The orange squares encompass the mating-type loci and indicate genome regions where the repeat contents are high and sequences are generally mating-type specific. (**B**) Model that explains the mating-type specificity of sequences within and flanking the mating-type loci. The absence of recombination within the mating-type loci and their flanking regions makes that transpositions within these regions become mating-type specific.

**Table 1.** Genome statistics of various smut genome assemblies.

| Species | *U. hordei* |  |  |  |  |  | *U. nuda* | *U. brachipodii-*<br>*distachyi*(27) | *U. tritici* | *U. lolicola* | *U. maydis*(25) | *S. reilianum*(37,<br>38) |
| --- | --- | --- | --- | --- | --- | --- | --- | --- | --- | --- | --- | --- |
| Strain | Uh359 | Uh805 | Uh811 | Uh818 | Uh1273 | Uh1278 | DE_29490 | UB2112 | Ut_3 | Us_530 | 521 | SRS1_H2-8 |
| Assembly size (Mb) | 27.0 | 25.8 | 26.2 | 26.2 | 27.2 | 26.6 | 21.4 | 20.4 | 20.4 | 20.8 | 19.7 | 18.5 |
| Contigs | 46 | 23 | 26 | 25 | 37 | 27 | 31 | 23 | 32 | 41 | 27 | 23 |
| BUSCOs (%) | 98.9 | 98.9 | 98.9 | 98.9 | 98.6 | 99.0 | 98.9 | 99.1 | 98.9 | 98.8 | 98.8 | 98.5 |
| Telomers* | 14 | 22 | 19 | 20 | 23 | 23 | 45 | 37 | 47 | 43 | 1 | 0 |
| Total repeats (%) | 38.2 | 35.3 | 36.4 | 36.5 | 38.9 | 36.0 | 22.6 | 17.0 | 16.4 | 8.9 | 4.6 | 3.6 |
| Class I TEs (kb)<sup>§</sup> | 5,625 | 4,611 | 4,985 | 4,897 | 5,663 | 4,940 | 1,786 | 672 | 739 | 102 | 199 | 5 |
| LTR (kb) | 5,272 | 4,326 | 4,615 | 4,549 | 5,208 | 4,607 | 1,537 | 463 | 462 | 9 | 185 | 5 |
| Gypsy (kb) | 2,066 | 1,688 | 1,679 | 1,653 | 2,225 | 1,873 | 484 | 144 | 127 | 3 | 4 | 3 |
| Copia (kb) | 2,732 | 2,331 | 2,561 | 2,554 | 2,587 | 2,531 | 1,1019 | 292 | 289 | 6 | 182 | 1 |
| Class II TEs (kb)<sup>§</sup> | 781 | 746 | 708 | 791 | 731 | 692 | 395 | 473 | 285 | 482 | 5 | 103 |

<sup>§</sup> Only repetitive sequences that were larger than 500 bp were classified.
\* “TAACCC” or “GGGTTA” repeats at the end of a contig

### *The U. hordei* secretome is activated upon plant colonization, whereas LTR-RTs are generally inactive

To study gene and LTR-RT expression, RNA-seq was done from from *U. hordei* grown in liquid medium, and in barley leaves at 3 days post infection. In total, 6229 of the 7704 (81%) predicted gene loci in Uh803 were expressed in either of the two *U. hordei* growth conditions, whereas only 27 of the 904 (3%) LTR-RTs displayed expression (Figure 2B).

**Figure 2.**
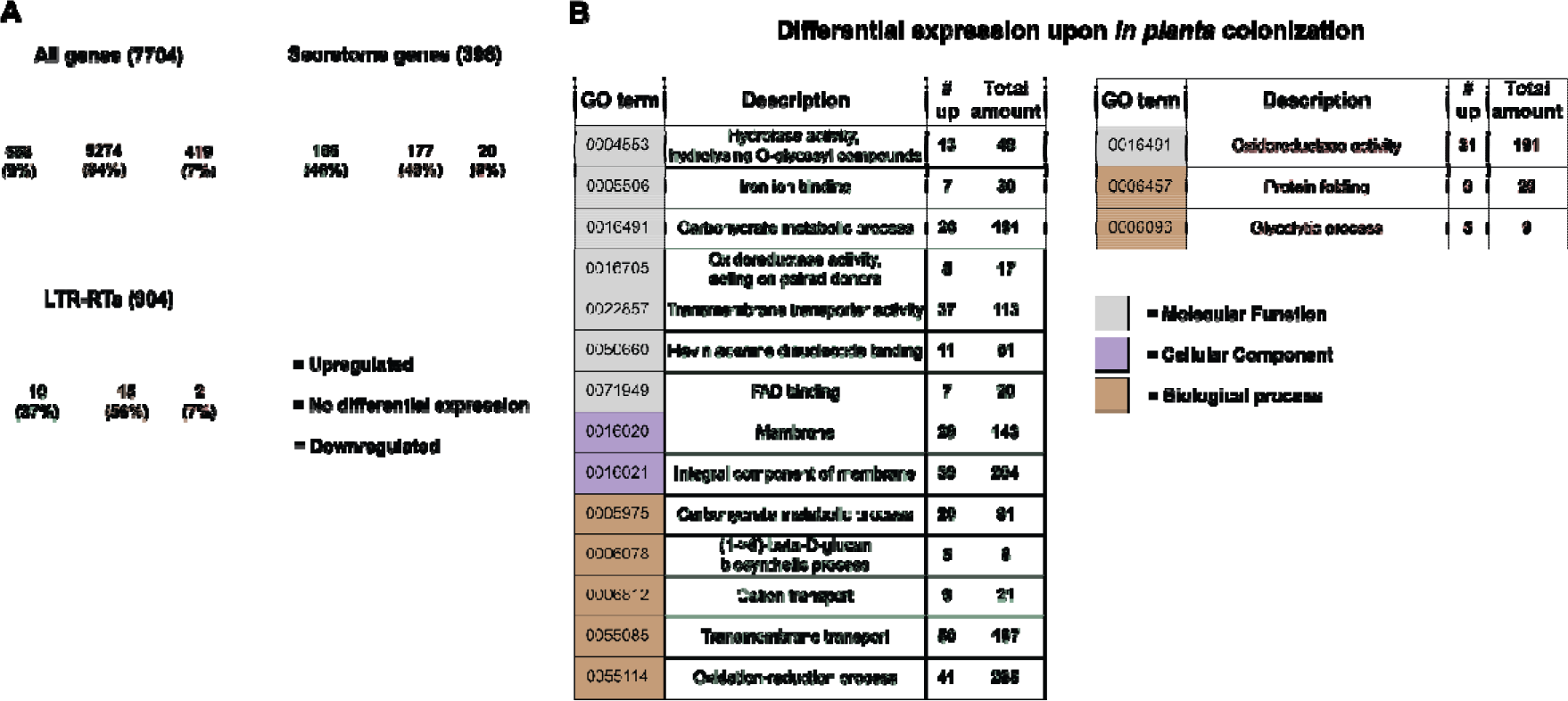
Differential expression of U. hordei loci upon plant colonization. (A) Comparison of U. hordei locus expression between growth in liquid culture medium and in planta. The numbers between brackets indicate how many genes and LTR-RTs have been annotated in the genome. The significance of differential expression was calculated using a threshold of log2-fold-change. (B) Gene Ontology (GO) term enrichments in differently regulated Ustilago hordei genes. In green and red are GO terms that are enriched in in plant up- and down-regulated genes, respectively. p-values were calculated with the Fisher’s exact test. For the whole figure significance was determined with a p-value < 0.01 and corrected for multiple-testing with the Benjamini- Hochberg method.

Moreover, only 7 of the expressed LTR-RTs displayed expression in more than half of their sequence. Of these 7, there was one *Copia* and one *Gypsy* LTR-RT that can be autonomous, as functional domains for aspartyl protease, reverse transcriptase and integrase could be identified. In conclusion, almost all LTR-RT sequences were inactive in the two tested environmental conditions. For the genes, 558 (9%) were upregulated *in planta*, whereas 419 (7%) were downregulated (Figure 2). Up- and downregulated genes were screened for Gene Ontology (GO) term enrichments, to see which biological processes are affected by plant colonization. In total, 14 and 3 GO terms were enriched in *in planta* up- and down-regulated genes, respectively (Figure 2). Generally, processes associated with the fungal membrane, including transmembrane transport were upregulated *in planta*. In correspondence with these results, 165 of the 558 genes upregulated *in planta* encode secreted proteins. Thus, 45% (165/369) of the expressed genes that encode secreted proteins were upregulated *in planta*, which is a significant enrichment (Fisher exact test, *p*-value = 1.87e-83) (Figure 2). In contrast, only 6% of the genes encoding a secreted protein were downregulated. Of these downregulated genes, 35% (7/20) was predicted to have a carbohydrate-active (CAZyme) function, whereas this was 18% (29/165) for *in planta* upregulated secretome genes. Thus, the *U. hordei* transmembrane transport system and secretome genes are strongly activated upon plant colonization, whereas hardly any LTR-RTs display expression. In total, 24% (median) of the 20kb flanking regions secretome genes consist of repeats, which is the same for non-secretome genes (t-test, *p*-value = 0.51) (Figure S2). Secretome genes upregulated *in planta* have a median of 21%, which is not significantly lower than non-secretome genes (t-test, *p*-value > 0.01). Thus, in contrast to some other filamentous plant pathogens (17), secretome genes are not especially associated with repeat-rich genome regions in *U. hordei*.

### Higher LTR-RT contents in genomes of smuts with a bipolar mating-type system

As LTR-RTs played a predominant role in the genome expansion of *U. hordei*, we also studied the impact of TE dynamics on the genome evolution of *U. hordei* sister species. We sequenced genomes of *Ustilago nuda*, *Ustilago tritici* and *Ustilago loliicola*, which are smut species that are close relatives of *U. hordei* and *U. brachipodii-distachyi* (29, 39). Assemblies of 21.4, 20.8 and 20.4 Mb were obtained in 31, 41 and 32 contigs for *U. nuda*, *U. loliicola* and *U. tritici*, respectively (Table 1). A phylogenetic tree was constructed, which included the newly sequenced species as well as *U. brachipodii-distachyi*, *U. maydis* and *S. reilianum* (Figure 3) (25, 27, 37, 38). *U. hordei, U. nuda*, *U. brachipodii-distachyi* and *U. tritici* cluster together with *U. loliicola* being the closest outgroup species. Within the cluster, *U. hordei* diverged the most recently from *U. nuda*, which also infects *Hordeum* species (Figure 3). Synteny between the different contigs was also investigated and the ancestral gene order reconstructed. *S. reilianum* and *U. loliicola* do not have inter-chromosomal rearrangement in comparison to their reconstructed last common ancestor (Figure 3). The *U. maydis* genome has one inter-chromosomal rearrangement with respect to its last common ancestor with *S. reilianum*. *U. hordei, U. nuda*, *U. brachipodii-distachyi* and *U. tritici* share one inter-chromosomal rearrangement that occurred after their divergence from *U. loliicola* (Figure 3). As previously reported, this rearrangement resulted in the mating-type polarity switch from tetrapolar to bipolar due to the linkage of the *a* and *b* mating-type loci (31). This inter-chromosomal rearrangement is the only one observed in the assemblies of *U. brachipodii- distachyi, U. nuda* and *U. hordei*, whereas the *U. tritici* assembly has one additional inter-chromosomal rearrangement. The smut species with a bipolar mating-type generally have a higher repeat content (16.4-38.9%), than the tetrapolar ones (3.6-8.9%) (Table 1). This increase in repeat content can largely be attributed to LTR-RT sequences, which comprise 4,326 kb in *U. hordei* (Uh805) in contrast to only 5 kb for *S. reilianum* (Table 1). Thus, repeats have increased after the polarity switch, mainly due to higher LTR-RT contents. Furthermore, repeat and the LTR-RT contents of smut genomes with a bipolar mating type positively correlate to mating-type loci sizes (*r* = 0.98, *p*-value = 0.02, using strain Uh805 for *U. hordei*), which ranges from 190 kb for *U. brachipodii-distachyi* to 560 kb for *U. hordei* (Figure 3, Table S1). In conclusion, the proliferation and/or retention of TEs seems to be an important determinant of the eventual size of mating-type loci.

**Figure 3:**
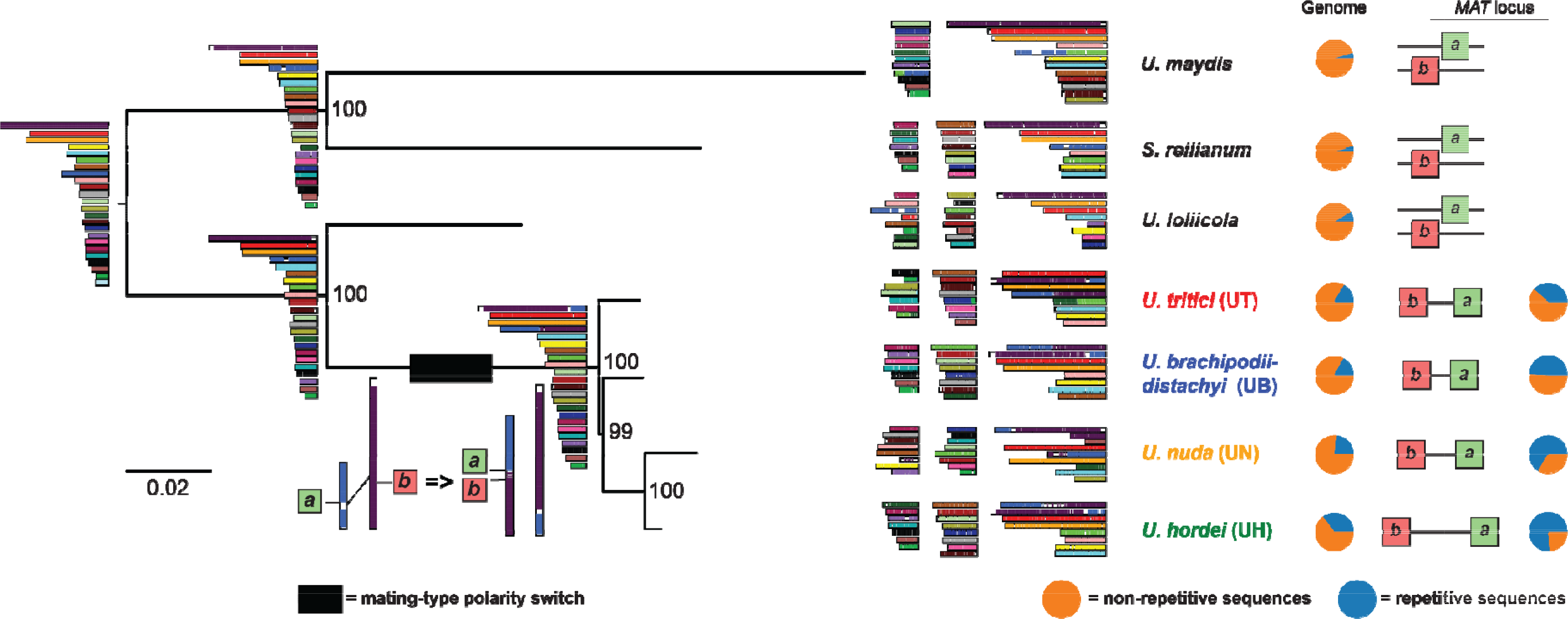
Genome evolution of smut species with bi-and tetrapolar mating systems. (**A**) Phylogenetic relationship between smut pathogens based on Benchmarking Universal Single-Copy Orthologs (BUSCOs). Phylogenetic relationship between newly and previously sequences smut species was constructed with the *Ustilago maydis*/*Sporisorium reilianum* branch as an outgroup. In total, 1,667 BUSCOs were used for tree construction. For *U. hordei*, strain Uh805 was used in the tree. The robustness of the trees was assessed using 100 bootstrap replicates. The colours of the contigs indicate the synteny with the ancestor contigs. The blue section of the circles indicate the repeat fraction that is present in the genome assembles and within the mating-type loci for species with a bipolar mating-type System.

### The timepoint of most recent LTR-RT proliferation differs between smut species

Although species with a bipolar mating system collectively encountered an increase in LTR-RT content, there are large interspecific differences as *U. hordei* has more than 9 times the amount of LTR-RT sequences than *U. tritici* (Table 1). To study the relative timepoint of the most recent LTR-RT proliferation, the nucleotide sequence identity distributions of the best reciprocal paralogous and orthologous LTR-RT sequences were calculated (Figure 4A). This was on the one hand done for the species with the highest LTR-RT contents, *U. hordei* and *U. nuda,* and on the other hand for *U. brachipodii-distachyi* and *U. tritici*. The distribution of the paralogous LTR-RTs in *U. brachipodii-distachyi* and *U. tritici* displayed two maxima, i.e. at 82-83% and at 88-90% (Figure 4A). The maximum of the orthologous LTR-RTs between *U. brachipodii-distachyi* and *U. tritici* was at 93%. Thus, orthologous LTR-RTs generally have a higher identity than paralogous ones, which indicates that LTR-RTs mainly proliferated before the last common ancestor of *U. brachipodii-distachyi* and *U. tritici* (Figure 4A). Orthologous LTR-RTs between *U. hordei* and *U. nuda* displayed a maximum at 89%, whereas for paralogous LTR-RTs a maximum at 93% was present for *U. nuda* and two maxima at 94 and 97% for *U. hordei* (Figure 4A). Thus, in contrast to *U. brachipodii- distachyi* and *U. tritici*, paralogous LTR-RTs generally have a higher identity than orthologous ones, which means that LTR-RTs continued to proliferated after the last common ancestor of *U. nuda* and *U. hordei*.

**Figure 4:**
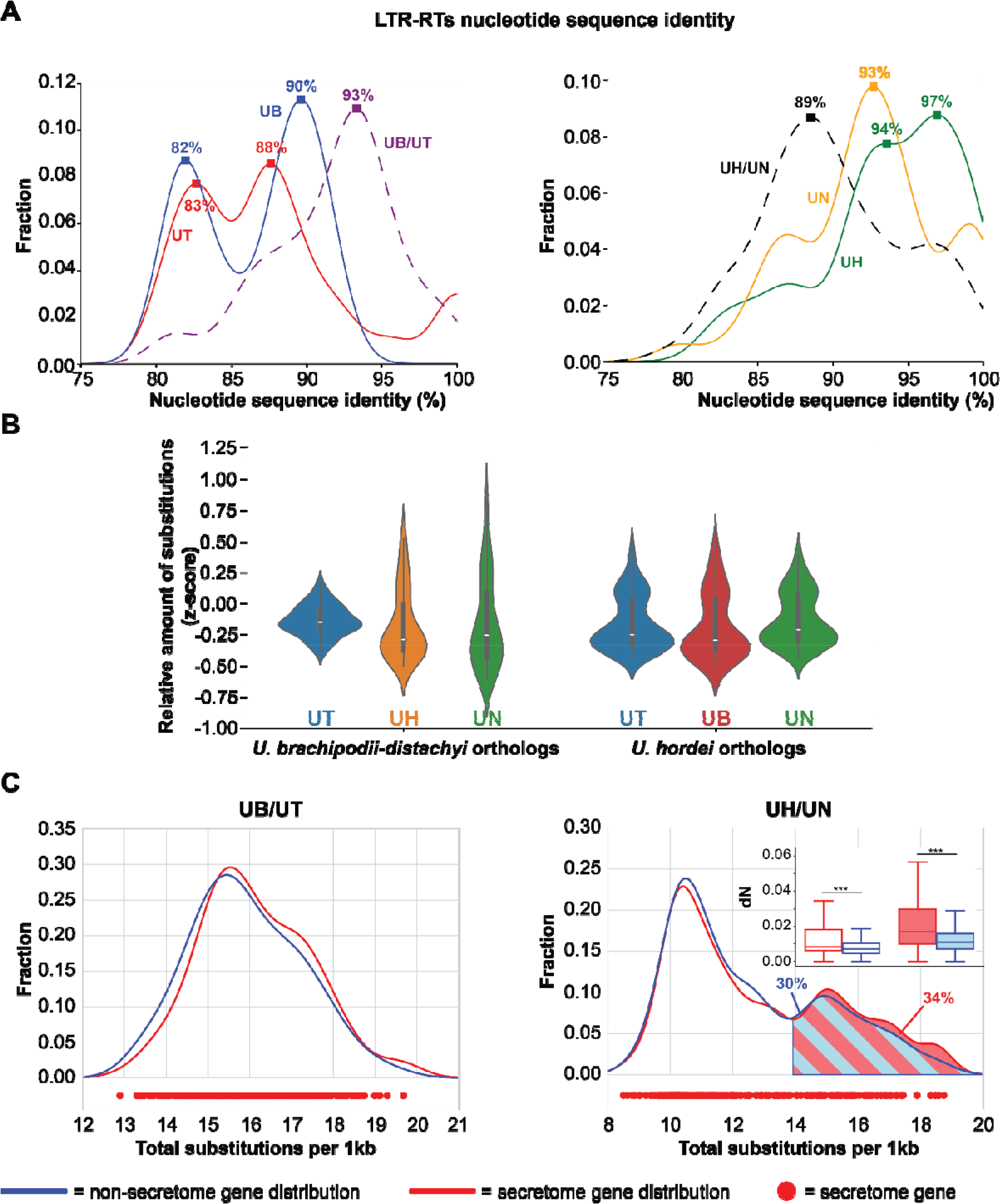
Interspecific comparison in long terminal repeat retrotransposon (LTR-RT) proliferation and local gene nucleotide substitution levels (A) Nucleotide sequence identity distribution of best reciprocal paralogous (full lines) and orthologous (striped lines) LTR-RT sequences. Squares on the lines display maxima with the corresponding sequence identity value. (B) The normalized sequence identity (z-score) was calculated for *U. brachipodii-distachyi* and *U. hordei* genes with orthologs of other bipolar mating-type species. The sequence identity was determined for non-overlapping sliding windows of 75 genes. (C) The distribution of the sequence divergence between *U. brachipodii-distachyi*/*U. tritici* and *U. hordei*/*U. nuda* ortholog windows (75 genes) are depicted for secretome and non-secretome genes in red and blue, respectively. The *U. hordei*/*U. nuda* distribution displays two peaks. The number of nonsynonymous substitutions per nonsynonymous site (dN) was compared between secretome and non-secretome genes for genes in the first and second distribution peak. Significance was determined with an unequal variance t-test. ***: *p*-value < 0.001.

### High nucleotide substitution levels affect secretome proteins

As TE-active genome regions have been associated with distinct nucleotide substitution regimes (11, 40, 41), we studied if different extents of LTR-RT fractions are associated with different nucleotide substitution regimes. We calculated the median number of substitutions between orthologs in windows of 75 genes. To ensure that genes are transcriptionally active, we only analysed *U. hordei* genes that displayed expression from here onwards. The variation in the normalized number of nucleotide substitutions (z-score) between *U. brachipodii- distachyi* and *U. tritici* ortholog windows is around 5.3 and 8.8 times less than *U. brachipodii-distachyi* ortholog windows with *U. hordei* and *U. nuda*, respectively (Figure 4B). In contrast, nucleotide substitutions of *U. hordei* ortholog windows with the other bipolar mating-type species display a more constant variation as the most varying ortholog windows (with *U. brachipodii-distachyi*) have only a 0.5 times higher variation than the least varying (with *U. nuda*) (Figure 4B). Thus, since their last common ancestor, gene nucleotide sequence divergence occurred more evenly across the genomes of *U. brachipodii-distachyi* and *U. tritici* than in *U. hordei* and *U. nuda*, where the divergence is more clustered. Correspondingly, substitutions between *U. brachipodii-distachyi* and *U. tritici* ortholog windows have a unimodal distribution, whereas the distribution between *U. hordei* and *U. nuda* have two distinct peaks (Figure 4C). For both comparisons, the distributions of secretome genes generally corresponds to that of non-secretome genes (Figure 4C). For *U. hordei*/*U. nuda* ortholog windows, the second peak in the distribution contains 30% of the non-secretome and 34% of the secretome genes, which is not significantly different (Fisher exact test*, p*-value = 0.10). Thus, high nucleotide substitution levels are not especially associated with secretome genes. Also for the second distribution peak, no GO terms enrichments could be found (*p*-value < 0.01). Furthermore, nucleotide substitution levels are negligibly positively correlated (Pearson’s r = 0.14 and *p*-value = 0.0026) with the fraction of species-specific genes (*U. hordei* genes without *U. maydis* ortholog) (Figure S3). In conclusion, genes in genome regions with high nucleotide substitution levels could not be associated with a particular function or more clear accessory nature. However, higher nucleotide substitution levels have a different impact on genes depending on their function. Substitutions that lead to amino acid alterations are more frequently fixed in secretome genes than in non-secretome genes (Figure 4C). The median number of nonsynonymous substitutions per nonsynonymous site (dN) for secretome genes is 18% higher than for non-secretome genes in the first peak of the *U. hordei* and *U. nuda* secretome distribution, whereas this is 55% for the second peak. Thus, the more flexible nature of secretome proteins makes that a higher nucleotide substitution rate speeds up their evolution.

### High substitution levels are association with high fractions of CG substitutions

We then analysed which type of substitutions (AC, AG, AT, CG, CT, GT) occur across the different substitution levels. The number of all substitution types are positively correlated with the number of total substitutions. Transitions (AG and CT substitutions) are responsible for 56% of the different substitution levels between *U. brachipodii-distachyi*/*U. tritici* ortholog windows (Figure 5A). In total, 27% of the variance can be attributed to CG substitutions, whereas the other transversions ranged from 4 to 7%. Similarly, for *U. hordei*/*U. nuda* ortholog windows, the number of all substitutions types display a positive correlation with the number of total substitutions. Here, CG substitutions are responsible for 47% of the variation in nucleotide substitution levels, whereas the contributions of other substitution types range from 5% (GT) to 16% (CT) (Figure 5A). The fraction of CG substitutions varies from 4% to 27% across the ortholog windows, whereas this is 3% to 16% for *U. brachipodii-distachyi*/*U. tritici* ortholog windows (Figure S4A). Correspondingly, the number of all substitution types in intergenetic regions are positively correlated with the total number of gene substitutions (Figure 5B). Similar to the coding regions, transitions contributed 52% to the intergenic substitution variation, whereas this was 23% for CG substitutions and 8-9% for the other transversion in *U. brachipodii-distachyi*/*U. tritici* ortholog windows. In contrast, *U. hordei*/*U. nuda* ortholog windows, transitions only contributed 40% to the variation of intergenic substitution levels (Figure 5B, S4B). All substitution types considered, CG displayed the highest variation and was responsible for 24% of the total nucleotide substitution variation. Although CG has, with 24%, the highest variation, this contrast with the 47% of coding regions. This discrepancy may be due to the difference in selection regime between coding and non-coding genome regions. The dN for every individual substitution type is positively correlated with the total substitution level. The correlation slope is the highest for CG substitutions, which is 3.5 times higher than for the second highest slope (AT). Similarly, the number of synonymous substitutions per synonymous site (dS) also has the steepest correlation slope for CG. However, this slope is only 1.5 times greater than the second highest slope (CT). In conclusion, *U. hordei* and *U. nuda* encountered more variation in their local nucleotide regimes than *U. brachipodii- distachyi* and *U. tritici*. For *U. hordei* and *U. nuda*, genome regions with higher nucleotide substitution levels encountered a relatively higher fraction of CG substitutions, which, after selection, is especially apparent in coding regions. Conceivably, different contributions of substitutions types impact codon frequencies and consequently amino acid compositions of proteins. Encoded proteins of genes that reside in genome regions with higher substitution levels are Cys, Gln, His, Leu richer and Asp, Gly, Phe, Val poorer than regions with lower substitution levels (Figure S5). Moreover, these specific amino acid tendencies have become more aggravated since the *U. hordei* divergence from *U. brachipodii-distachyi* (Figure S5).

**Figure 5:**
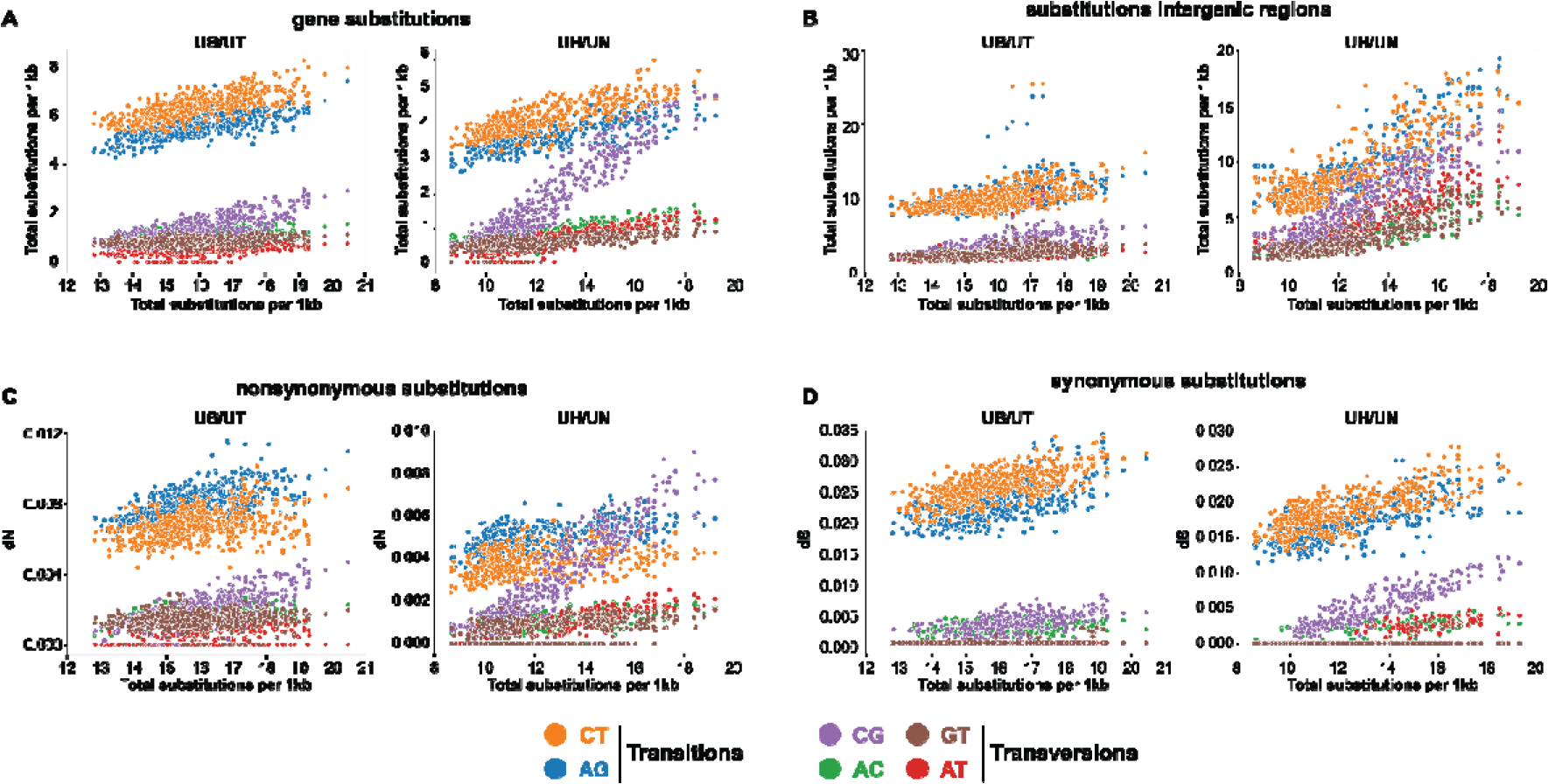
Comparison of nucleotide substitution regimes for U. brachipodii-distachyi/U. tritici and U. hordei/U. nuda ortholog windows. The nucleotide substitutions were calculated for windows of 75 genes with a sliding step of 10. The x-axis consistently displays the total substitutions per 1 kb for these windows. (A) The y-axis depicts the median number of every substation type (CT, AG, CG, AC, GT, AT) of ortholog windows. (B) The y-axis depicts the median number of every substation type for the intergenetic regions of ortholog windows. (C) The y-axis depicts the median fraction of nonsynonymous substitutions per nonsynonymous site (dN) for every substitution type in ortholog windows. (D) The y-axis depicts the median fraction of synonymous substitutions per synonymous site (dS) for every substitution type in ortholog windows.

### High local nucleotide substitution levels are associated with LTR-RT proliferation

As higher nucleotide substitution levels with distinct substitution patterns occur in *U. hordei* and *U. nuda*, which are species with more recent LTR-RT proliferations than *U. brachipodii- distachyi* and *U. tritici*, we looked for a direct association with LTR-RTs. The median distance of *U. hordei* genes to their closest LTR-RT is significantly, negatively correlated to the median substitution level (with *U. nuda* orthologs) of ortholog windows (Pearson’s r = - 0.27, *p*-value = 4.74e-9) (Figure 6A). A correlation coefficient of -0.27 points towards a weak correlation. In conclusion, genes in genome regions with higher nucleotide substitution levels generally reside closer to LTR-RTs.

**Figure 6:**
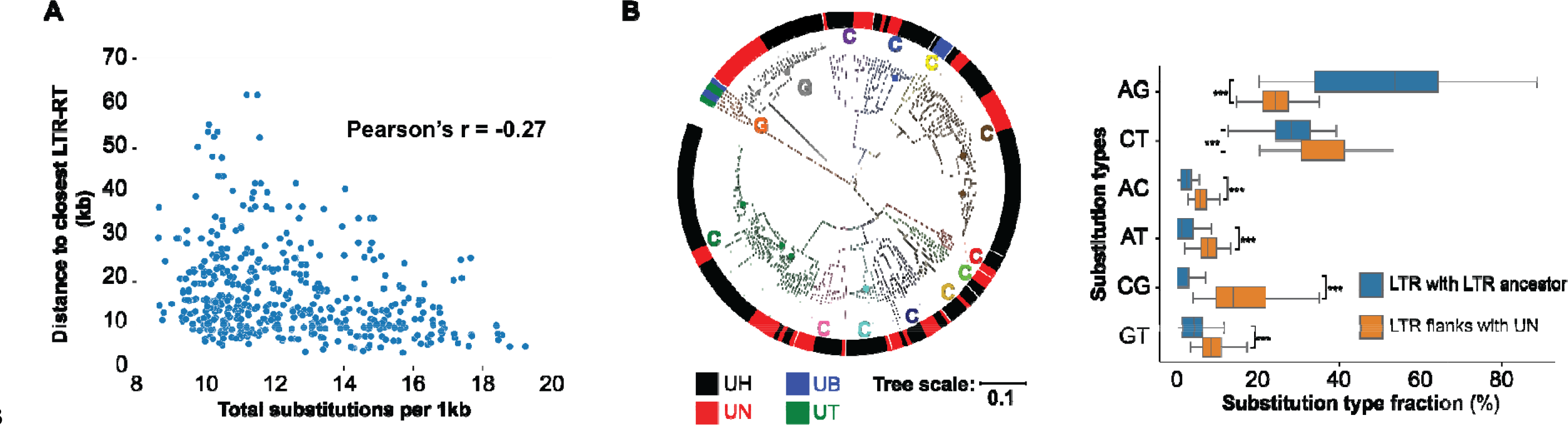
High local nucleotide substitution levels are associated with long terminal repeat retrotransposons (LTR-RTs). (364 A) The relation between the median number of nucleotide substitutions and the median distance between U. hordei genes and the closest LTR-RT for ortholog windows of 75 genes with a sliding step of 10. (B) In total, 252 LTR-RTs are included in the phylogenetic tree and their species of origin is indicated by the outer band colour (UH = U. hordei, UN = U. brachipodiidistachyi distachyi, UT = U. tritici). LTR-RTs families are indicated by the circle segments in different colour. Families indicated with “G” are gypsy-type families and “C” are copiatype type families. For recently proliferated U. hordei LTR-RTs, the fractions of the different substitution types were determined with their LTR-RT ancestor that is indicated with a circle on the phylogenetic tree. Substitution fractions of the 20 kb flanking regions (40 kb in total) of the LTR-RT with U. nuda, excluding repetitive sequences, were also determined. Significant differences between LTR and flanking regions were determined for every substitution type individually with an unequal variance t-test. ***: p-value <0.001.

To study the LTR-RT nucleotide substitution regime, we constructed ancestor LTR- RT sequences of LTR-RT families, using the convention that TE family members share at least 80% sequence identity in at least 80% of their sequence with one other family member (19). To facilitate the sequence alignment and ancestor sequence construction, we only took a subset of the LTR-RTs and excluded the terminal repetitive sequences (more details in Material and Methods). In total, ancestors of 13 LTR-RT families were reconstructed using 252 LTR-RT sequences (Figure 6B). We then determined clades in the phylogenetic tree that solely consists of very similar *U. hordei* LTR-RTs and, thus, recently proliferated in *U. hordei*. In relation to their ancestor sequences, LTR-RTs substitutions consisted 91% (median) of transitions (Figure S6). In contrast, nucleotide substitutions of their 20kb flanking regions (excluding repetitive sequences) consisted 62% of transitions (compared to. *U. nuda*). Here, CG comprised the highest fraction of transversions with a median of 14% of the total substitutions (Figure 6). In contrast, only 1% of the substitutions between LTR-RTs and their ancestors were CG. In conclusion, LTR-RTs are not subjected to the nucleotide substitution regime with a high fraction of CG substitutions.

## Discussion

Nucleotide substitution rates are unevenly distributed across genomes and can be influenced by numerous factors, including neighbouring nucleotides, recombination frequencies and TE activity (41–43). Nucleotide divergence in *U. hordei* and *U. nuda* occurred more clustered in their genomes compared to *U. brachipodii-distachyi* and *U. tritici* (Figure 4B-C). These differences in regional substitution rates can be directly or indirectly caused by distinct LTR-RT dynamics, as *U. hordei* and *U. nuda* encountered a more recent LTR-RT proliferation than *U. brachipodii-distachyi* and *U. tritici* (Figure 4A, Table 1). Moreover, another association between LTR-RTs and nucleotide substitution rates was found, as gene nucleotide substitution levels are weakly, negatively correlated to the distance of the closest LTR-RT in *U. hordei* (Figure 6A). Conceivably, the purge of LTR-RTs from the genome impacts this correlation considerably, as purged LTR-RTs cannot be detected, but may have had an impact on the local nucleotide substitution regime. High nucleotide substitution levels are accompanied with a high fraction of CG substitutions (Figure 5,S4). A relatively high fraction of CG substitutions is found in the flanking regions of recently proliferated LTR-RTs, but not for LTR-RTs themselves (Figure 6B). A mechanism to how LTR-RTs may impact local nucleotide substitution regimes remains elusive. The relation might be indirectly, and caused by different epigenetic regimes in the genome (44). Distinct methylation and/or histon modification patterns may occur in LTR-RT-rich genome regions, which leads to a more erroneous DNA polymerase with high CG substitutions. However, LTR-RTs themselves are not subjected to a high fraction of CG substitutions. Possibly, DNA methylation may specifically target LTR-RT sequences, which cause a distinct nucleotide substitution regime that is different from the LTR-RT flanking regions. Alternatively, the distinct nucleotide substitution regime may not have an epigenetic origin and originates from a more erroneous DNA polymerisation of the single stranded LTR-RT flanking regions during LTR-RT insertion. This mechanism has been previously suggested in rice, where higher nucleotide substitutions levels occur close to TE insertion sites (41). TE insertion causes cuts in the host DNA, which are then ligated by the host (45, 46). However, the cut host DNA might become a target for 3’->5’ exonuclease resulting in a segment of single-stranded DNA (41). The complementary strand of this stretch of DNA would then be synthesized by a replication complex with lower DNA polymerase fidelity and mismatch repair. This hypothesis could explain why the nucleotide substitution regime with high CG fractions affect LTR-RT neighbouring regions, but not LTR-RTs themselves.

Higher levels of nucleotide substitutions impact the evolution of the genes that reside in the affected genome regions. Particularly the occurrence of nonsynonymous CG substitutions strongly increases with higher substitution levels (Figure 5C). These shifts in nucleotide substitution regime change the amino acid composition of proteins (Figure S5). High nucleotide substitution levels especially lead to amino acid alteration in secretome genes, as they are more flexible to amino acid changes than other genes (Figure 4C). Although the effect of nucleotide substitutions affected secretome genes more, enrichments of particular gene functions could not be found for genome regions with high nucleotide substitution levels. Thus, the high substitution levels are not in line with the two-speed genome model (18, 24), as they do not specifically affect genome regions that are rich in secretome genes, which include effector gene candidates. More generally, repeat content were not more frequently found in the proximity of secretome genes compared to other genes (Figure S2). The specificity and the universality of the two-speed genome model for filamentous plant pathogens has recently been contested (47, 48). More plant pathogens have been reported where effector candidates do not especially reside in gene-poor/repeat-rich regions, such as the leaf spot pathogen *Ramularia collo-cygni* on barley, the earlier mentioned *P. striiformis* f. sp. *tritici* and the barley powdery mildew pathogen *Blumeria grami*nis f. sp. *hordei* (49–51).

LTR-RTs are mainly responsible for the *U. hordei* genome expansion (Table 1). The expansion occurred especially in the mating-type locus that increased almost three times in size in comparison to *U. brachipodii-distachyi* (Table S1). The lack of the purifying recombination ability in this genome region can be the reason why LTR-RTs especially accumulated in the mating-type and flanking genome regions (28, 32). Conceivably, this process is reinforced by the increasing presence of repetitive sequences as the transposition into a repeat-rich genome region is less likely to have a severe fitness cost than a transposition into repeat-poor regions. Furthermore, the co-occurrence of high LTR-RTs genome contents and the switch in mating-type organization from tetra- to bipolar may indicate that mating-type polarity impacts LTR-RT proliferation and/or retention (52). In case of biallelic *a* and *b* loci, the switch from a tetra-to bipolarity results in a basidiospore compatibility change from 25 to 50%. Consequently, it takes a tetrapolar smut on average longer to find a mating type than a bipolar smut. This longer time might increase the opportunity to mate with spores from a different offspring and, thus, increase outcrossing. The higher outcrossing rate for tetra- compared to bipolar smuts is even more pronounced when multiallelism exists for the *a* and *b* loci (53). Multiallelism increases the compatibility on population level, whereas compatibility within the same offspring remains 25 and 50% for tetra- and bipolar smuts, respectively. Lower levels of outcrossing reduce the purifying recombination ability of smuts, which may be the reason why LTRs could be retained for longer and proliferate to a further extent in bipolar smuts (28, 52).

TEs are import drivers of genome evolution as they directly cause mutagenesis through their transpositions and indirectly increase the change of non-homologous recombination due to their repetitive nature (12). LTR-RT proliferation in *U. hordei* indicates that TE activity may also influence local nucleotide substitution regimes and increase the substitution levels in the genome regions where they insert. Consequently, genes in the proximity of these insertion sites encounter more nonsynonymous substitutions and thus evolve faster (Figure 5C). Fast gene evolution may be advantageous under stressful condition, when TEs are typically more active or change their activity (54, 55).

## Material and Methods

### Genome sequencing and assembly

Genomic DNA from all smut species was isolated using a MasterPure™ Complete DNA&RNA Purification Kit (Epicentre®, Illumina®, Madison, Wisconsin, USA) according to the manufacturer’s instructions. Long *U. hordei* reads were obtained with the Oxford Nanopore MinION device. The genomes of six *U. hordei* strains were sequenced: Uh359, Uh805, Uh811, Uh818, Uh121 and Uh122 (10). The library was prepared according the Oxford Nanopore Technology (ONT) protocol for native barcoding genomic DNA (EXP- NBD104 and SQK-LSK109). Three *U. hordei* strains were multiplexed for every run. The prepared library was loaded on an R9.4.1 Flow Cell. ONT reads were base-called, filtered (default value) and barcodes were trimmed with the Guppy Basecalling Software v3.5.1 of ONT. Paired-end *U. hordei* 150 bp reads were obtained with the Illumina HiSeq 4000 device. Library preparation (500bp insert size) and sequencing were performed by the BGI Group (Beijing, China). Paired-end *U. hordei* reads were filtered using Trimmomatic v0.39 with the settings “LEADING:3 TRAILING:3 SLIDINGWINDOW:4:15 MINLEN:100”, only reads that remained paired after filtering were used in the assembly (56). In total, 3.2-4.5 Gb of filtered paired-end reads and 1.5-6.5 Gb of filtered Nanopore reads were used for assembly. An initial assembly was obtained by using the “ONT assembly and Illumina polishing pipeline” (https://github.com/nanoporetech/ont-assembly-polish). The assembly was further upgraded using the FinisherSC script (57) Mitochondrial contigs were removed from the assembly and were not used for any analysis. Additionally, small contigs were removed that contained a paired-end read coverage lower than 50% of the genome-wide average.

Long *U. nuda*, *U. loliicola* and *U. tritici* reads were obtained through Single Molecular Real-Time (SMRT) Sequencing using the PacBio Sequel system. A total of 6.3-9.7 Gb of raw long reads were obtained for the different species. The initial assembly was obtained using the Canu assembler and was further upgraded with the FinisherSC script (57, 58). Mitochondrial contigs were remove from the assembly and were not used for further analysis.

The quality of genome assemblies was assessed by screening the presences of BUSCOs using the BUSCO software version 5.0.0 with the database “basidiomycota_odb10” (59).

### Transposable element annotation and classification

The smut genome assemblies were scavenged for repetitive sequences in order to construct a repeat library for repeat annotation. Helitron TEs were identified using the EAHelitron script (60). LTR-RTs were identified using LTRharvest (61). Miniature inverted-repeat TEs were identified with MITE Tracker (62). Short interspersed nuclear elements were identified with the SINE-scan tool (63). Finally, RepeatModeler (v1.0.11) was also used for *de novo* repeat identification. These repeats were than combined with the repeat library from RepBase (release20170127) (64). The CD-HIT-EST tool under default settings was used to remove redundancy in the constructed library (65). RepeatMasker (v4.0.9) was then used to annotate the repeats to specific genome locations. The annotated repeat sequences were filtered on size and only sequences larger than 500bp were retained. Furthermore, repeats that were nested or had more than 50% overlap with other repeats were removed from the library. In case two repeats had reciprocally 50% overlap was the longest repeat retained. Repeats were classified into different TE orders using the PASTEC tool using PiRATE-Galaxy (66, 67).

### *U. hordei* RNA sequencing and expression analysis

Total RNA from *U. hordei* strain 4857-4 strain grown axenically and *in planta* was extracted for three biological replicates. For the axenic samples, *U. hordei* was grown in YEPS light (0.4% yeast extract, 0.4% peptone, and 2% saccharose) liquid medium at 22°C with 200 rpm shaking till OD:1.0. For the *in planta* samples, Golden Promise barley cultivar was grown in a greenhouse at 70% relative humidity, at 22°C during the day and the night; with a light/dark regime of 15/9 hrs and 100 Watt/m^2^ supplemental light when the sunlight influx intensity was less than 150 Watt/m^2^. Barley plants were infected with *U. hordei* through needle injection as previously described (68) and samples were harvested 3dpi. Here, the third leaves of the *U. hordei* infected barley plants were collected by cutting 1 cm below the injection needle sites. Leaf samples were then frozen in liquid nitrogen and grinded using a mortar and pestle under constant liquid nitrogen. The total RNA was isolated by using the TRIzol® extraction method (Invitrogen; Karlsruhe, Germany) according to the manufacturer’s instructions. Subsequently, total RNA samples were treated with Turbo DNA-Free™ Kit (Ambion/Applied Biosystems; Darmstadt, Germany) to remove any DNA contamination according to the manufacturer’s instructions. Total RNA was then sent to for library preparation and sequencing to Novogene (Beijing, China). Libraries (250-300 bp insert size) were loaded on Illumina NovaSeq6000 System for 150bp paired-end sequencing using a S4 flowcell.

In total, 5.1-8.4 and 36.0-45.2 Gb of raw reads were obtained for the samples grown in liquid medium and *in planta*, respectively. The reads were filtered using the Trinity software (v2.9.1) option trimmomatic under the standard settings (69). The reads were then mapped to the reference genome using Bowtie 2 (v2.3.5.1) with the first 15 nucleotides on the 5’-end of the reads being trimmed due to inferior quality (70). The reads were mapped onto a combined file of the *U. hordei* strain Uh114 genome assembly and the *Hordeum vulgare* (IBSC_v2) (71) genome assembly. Reads were counted to the *U. hordei* loci using the R package Rsubread (v1.34.7) (72). Here the default minimum mapping quality score of 0 was used, to include reads that would have multiple best mapping locations. For the gene loci, reads were counted that were mapped to the predicted coding regions. For the LTR-RT loci, reads were only counted that mapped within LTR-RT loci, excluding the reads that mapped onto the 10% of either edge of the locus. Loci were considered expressed if they had more than one count per million in at least two of the six samples (three replicates of two treatments). Significant differential expression of a locus was calculated using the R package edgeR (v3.26.8), using the function “decideTestsDGE” (73). Here, a threshold of log2 fold change of 1 was used and differential expression was determent using a p-value < 0.01 with Benjamini-Hochberg correction

### Gene annotation

*U. hordei* genomes were annotated using the BRAKER v2.1.4 pipeline with RNA-Seq and protein supported training with the options “--softmasking” and “--fungus” enabled (74). RNA-seq reads from *U. hordei* grown in axenic culture and *in planta* (all replicates) were mapped to the assemblies using TopHat v2.1.1 (75). Protein predictions from numerous Ustilaginales species were used to guide the annotation, i.e. *Anthracocystis flocculosa*, *Melanopsichium pennsylvanicum*, *Moesziomyces antarcticus*, *S. reilianum*, *U. brachipodii- distachyi*, *U. hordei*, *U. maydis* (25–28, 76–78). *U. nuda* and *U. tritici* genomes were also annotated with the BRAKER v2.1.4 pipeline, but no RNA-seq data was used to guide the annotation. The option “--fungus” was enabled and the previously published protein files of the following species were used for protein supported training: *M. pennsylvanicum*, *S. reilianum*, *U. brachipodii-distachyi* and *U. maydis* (25–27, 76). Our annotation of *U. hordei* Uh805 was also included to train the annotation software. The *U. brachipodii-distachyi* and *U. maydis* genomes were previously annotated and this annotation was used for analysis (25, 27). Predicted genes that included an internal stop codon or did not start with a methionine were removed.

Secreted proteins are proteins with a predicted signal peptide using SignalP version 5.0 (79) and the absence of a transmembrane domain predicted with TMHMM2.0c in the protein sequence excluding the signal peptide (80). Gene Ontology (GO) terms were annotated to the *U. hordei* strain Uh114 protein prediction using InterProScan (v5.42-78.0) (81). Significance of GO term enrichments in a subset of genes were calculated with a Fisher exact test with the alternative hypothesis being one-sided (greater). The significance values of the multiple enrichments were corrected according Benjamini and Hochberg (82). Carbohydrate-Active enzymes (CAZymes) were annotated using the dbCAN2 meta server (83, 84). A protein was considered a CAZyme if at least two of the three tools (HMMER,DIAMOND and Hotpep) predicted a CAZyme function.

### Comparative genomic analyses

Phylogenetic trees were constructed based on BUSCOs from the database “basidiomycota_odb10” that are present without paralog in all members of the tree (59). For every gene, the encoded protein sequences were aligned using MAFFT (v7.464) option “-- auto” (85). These aligned protein sequences were then concatenated for every species and used for tree construction using RAxML (v8.2.11) with substitution model “PROTGAMMAWAG” and 100 bootstraps (86). Here, protein sequences that were present in at least 60% of the tree members were excluded for tree construction.

Synteny block between the smut genome assemblies of were identified with SynChro with DeltaRBH = 3 (87, 88). The genome assembly of the epiphytic yeast *Moesziomyces bullatus* ex *Albugo* was included in this analysis to use as an outgroup (89). The ancestral chromosome gene order was constructed with AnChro with Deltá = 3 and Deltá = 3 (88, 90). Inter-chromosomal rearrangements, i.e. translocations of two blocks, were identified with ReChro Delta = 10 (88, 90). No inter-chromosomal rearrangements in *U. nuda* could be automatically detected by ReChro. Here, the inter-chromosomal rearrangement that lead to a mating-type polarity switch was manually determined.

To determine the specificity of *MAT* locus sequences, absent/present polymorphisms between *U. hordei* strains were determined with NUCmer (version 3.1) from the MUMmer package with the option “--maxmatch” (91). From the same package, delta-filter with the option “-1” was used to find the one-to-one alignments.

### LTR-RT evolution

To know the sequence identity distribution, the best orthologous and paralogous LTR-RTs were identified using blastn (v2.2.31+) (92). LTR-RTs that did not belong to an LTR-RT family of multiple members, were excluded from the analysis. Members of the same LTR-RT family share at least 80% sequence identity in at least 80% of their sequence with at least one other member (19). Orthologous or paralogous LTR-RTs that have reciprocally the highest bit-score were used for analysis. The nucleotide identity distribution of these orthologous and paralogous LTR-RTs was constructed using Gaussian Kernel Density Estimation with a kernel bandwidth of 1.5.

To reconstruct the ancestor LTR-RTs, a subset LTR-RTs were used. LTR-RTs were included that were larger than 3 kb and smaller than 15 kb. Furthermore, repetitive sequences within the LTR-RT (>50 bp) were indicated using blastn (v2.2.31+) and removed from the sequence (92). The region between the repeats were then used for ancestor construction if this region was larger than 500 bp. Here, bedtools (v.2.29.2) function “getfasta” was used (93). Open reading frames (ORFs) and there encoding amino acid sequence of were determined with esl-translate (-l 50) as part of the Easel (v0.46) package. Functional domain within these amino acid sequences were determined with pfam_scan.pl (-e_seq 0.01) using the Pfam database version 32.0 (94). Only sequences were included in the ancestor construction if they had at least 3 different Pfam domains from the following domains: PF00078, PF00665, PF03732, PF07727, PF08284, PF13975, PF13976, PF14223, PF17917, PF17919 and PF17921. All of these predicted Pfam domain had to located on the same nucleotide strand in order to be used for ancestor construction. These sequences were then grouped in families, according to the definition that family members share at least 80% sequence identity in at least 80% of their sequence with at least one other member (19). Families were classified in *Copia* or *Gypsy* using the tool LTRclassifier (95). Ancestors were constructed using prank (v.170427) with the options “-showall” and “-F”. Nucleotide substitutions between LTR-RTs and their constructed ancestor were then determined after they were aligned using MAFFT (v7.464) options “--auto” (85).

### Gene divergence

One-to-one orthologs and homologs between *U. hordei* strains were found using the SiLiX (v.1.2.10-p1) software with the setting of at least 35% identity and 40% overlap (96). Homolog groups consisting of two members, each one of a different strain/species, were considered one-to-one homologs. Nucleotide substitutions for orthologs were identified after orthologs were aligned using MAFFT (v7.464) options “--auto” (85). Synonymous and nonsynonymous substitutions between orthologs were identified using SNAP (97). The nucleotide substitution level distributions were constructed using Gaussian Kernel Density Estimation with a kernel bandwidth of 0.5.

### Data accession

Raw RNAseq reads and genome assemblies are deposited at NCBI under the BioProject PRJNA698760.

## Acknowledgements

This work has been supported by the Alexander von Humboldt Foundation, the European Research Council (ERC 2017 COG 771035, conVIRgens), the Cluster of Excellence on Plant Sciences (CEPLAS; Germany’s Excellence Strategy–EXC 2048/1 – Project ID: 390686111) and the University of Cologne. We also thank the Regional Computing Centre of Cologne (RRZK) for access to the Cologne High Efficient Operating Platform for Science (CHEOPS). We thank Guus Bakkeren for sharing the *U. hordei* strains with us and critically reading the manuscript. We thank Karl-Josef Müller for the isolation and sharing of the *U. nuda* and *U. tritici* strain with us.

## SUPPLEMENTAL MATERIAL

Table S1. Characteristics of the mating-type loci of smut species with a bipolar mating-type system.

Table S2. Transposable elements annotation in various smut genome assemblies.

Figure S1. **Phylogenetic relationship between *Ustilago hordei* lineages based on Benchmarking Universal Single-Copy Orthologs (BUSCOs).** In total, 1,692 BUSCOs were used for tree construction. Homologous BUSCO protein sequences were aligned using MAFFT and then concatenated for tree construction using RAxML with substitution model “PROTGAMMAWAG”. *U. nuda* was used as on outgroup species to root the tree. The robustness of the trees was assessed using 100 bootstrap replicates.

Figure S2. **Comparison of repeat content of gene flanking regions between expressed secretome genes and other expressed genes.** Upregulated means a significantly higher expression *in planta* compared to growth in axenic culture. In total, 20 kb sequences on each side of the genes (40 kb in total) were considered as flanking regions. Significant differences were calculated with a two-sided T-test. No significant differences with *p*-value < 0.01 were found.

Figure S3. **Correlation between median nucleotide substitution level and fraction *U. hordei* (UH) specific genes for ortholog windows.** Ortholog windows of 75 UH genes with a sliding step of 10 were used to determine the number of substitutions with *U. nuda*. UH specific genes do not have an ortholog in *U. maydis*.

Figure S4. Comparison of nucleotide substitution type fractions for *U. brachipodii- distachyi*/*U. tritici* and *U. hordei*/*U. nuda* ortholog windows. The nucleotide substitutions were calculated for windows of 75 genes with a sliding step of 10. The x-axis consistently displays the total substitutions per 1 kb for these windows. (A) The y-axis depicts the fraction of every substation type (CT, AG, CG, AC, GT, AT) of ortholog windows. (B) The y-axis depicts the fraction of every substation type for the intergenetic regions of ortholog windows.

Figure S5. **Correlations between the nucleotide substitution levels and the encoded amino acid composition of genes.** Correlations were calculated for windows of 75 *U. hordei* genes with a sliding step of 10. Significant correlations, with *p*-value < 0.01, are indicated by a black edge around the square. (**A**) Correlations between amino acid compositions of encoded *U. hordei* proteins and the number of nucleotide substitutions with *U. nuda*. (**B**) Correlations between amino acid alternations for encoded *U. hordei* proteins and the number of nucleotide substitutions using *U. nuda* and *U. brachipodii-distachyi* orthologs as comparison.

Figure S6. Fractions of transitions and transversion of recently proliferated *U. hordei* long terminal repeat retrotransposons (LTR-RTs) and their flanking regions. The fraction of transitions and transversions between recently proliferated *U. hordei* LTR-RTs and their ancestors were determined. Fractions of the 20 kb flanking regions (40 kb in total) of the LTR-RT with *U. nuda* (UN) were also determined. Significant differences between LTR and flanking regions were determined for transitions and transversions separately with an unequal variance t-test. ***: *p*-value < 0.001.

## Notes

### Competing Interest Statement

The authors have declared no competing interest.

