## Supplementary material for "High nucleotide substitution rates associated with retrotransposon proliferation drive dynamic secretome evolution in smut pathogens": Figure S1

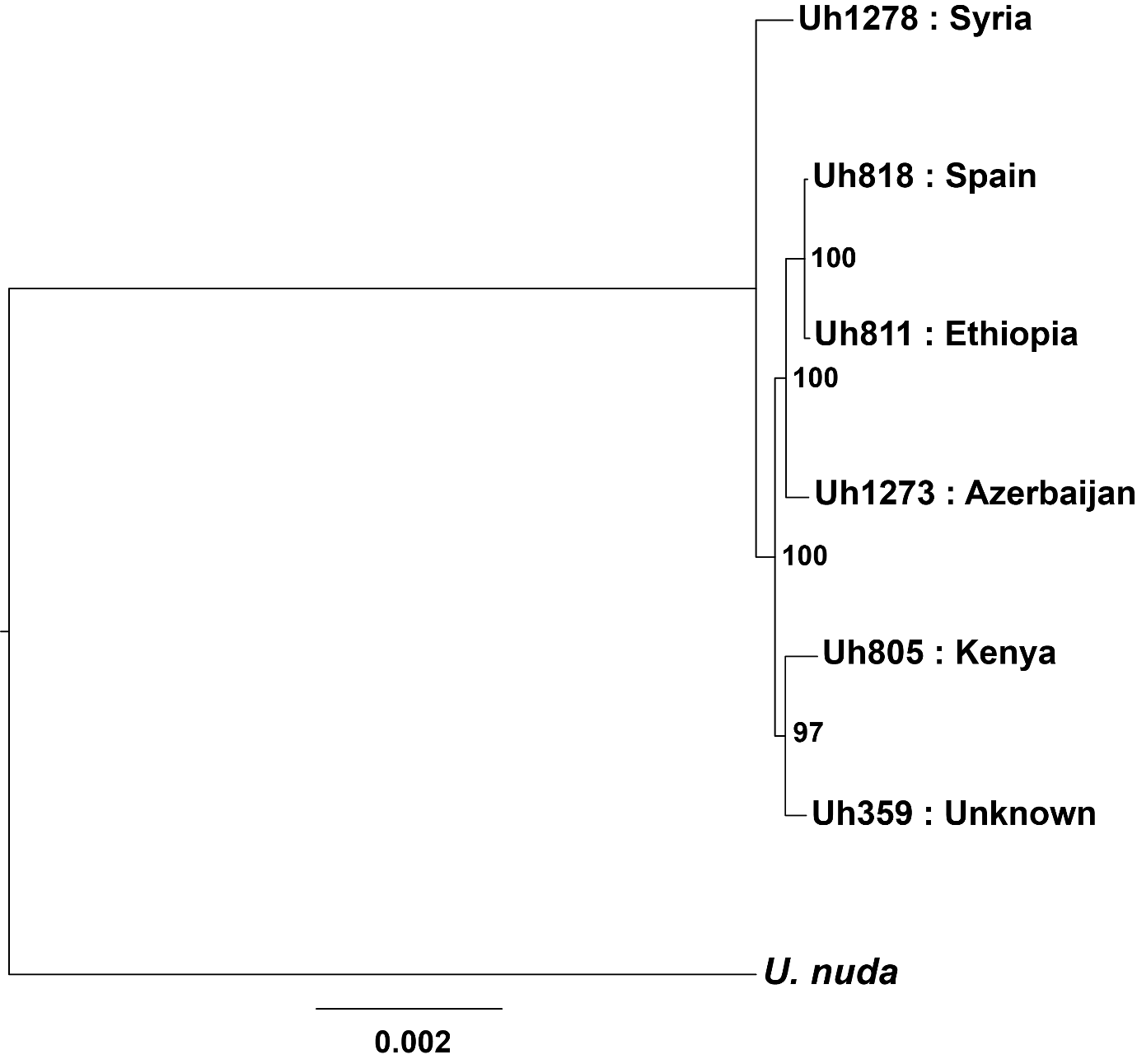

**Figure S1 Phylogenetic relationship between *Ustilago hordei* lineages based on Benchmarking Universal Single-Copy Orthologs (BUSCOs).** In total, 1,692 BUSCOs were used for tree construction. Homologous BUSCO protein sequences were aligned using MAFFT and then concatenated for tree construction using RAxML with substitution model “PROTGAMMAWAG”. *U. nuda* was used as on outgroup species to root the tree. The robustness of the trees was assessed using 100 bootstrap replicates.
