## Supplementary material for "High nucleotide substitution rates associated with retrotransposon proliferation drive dynamic secretome evolution in smut pathogens": Figure S2

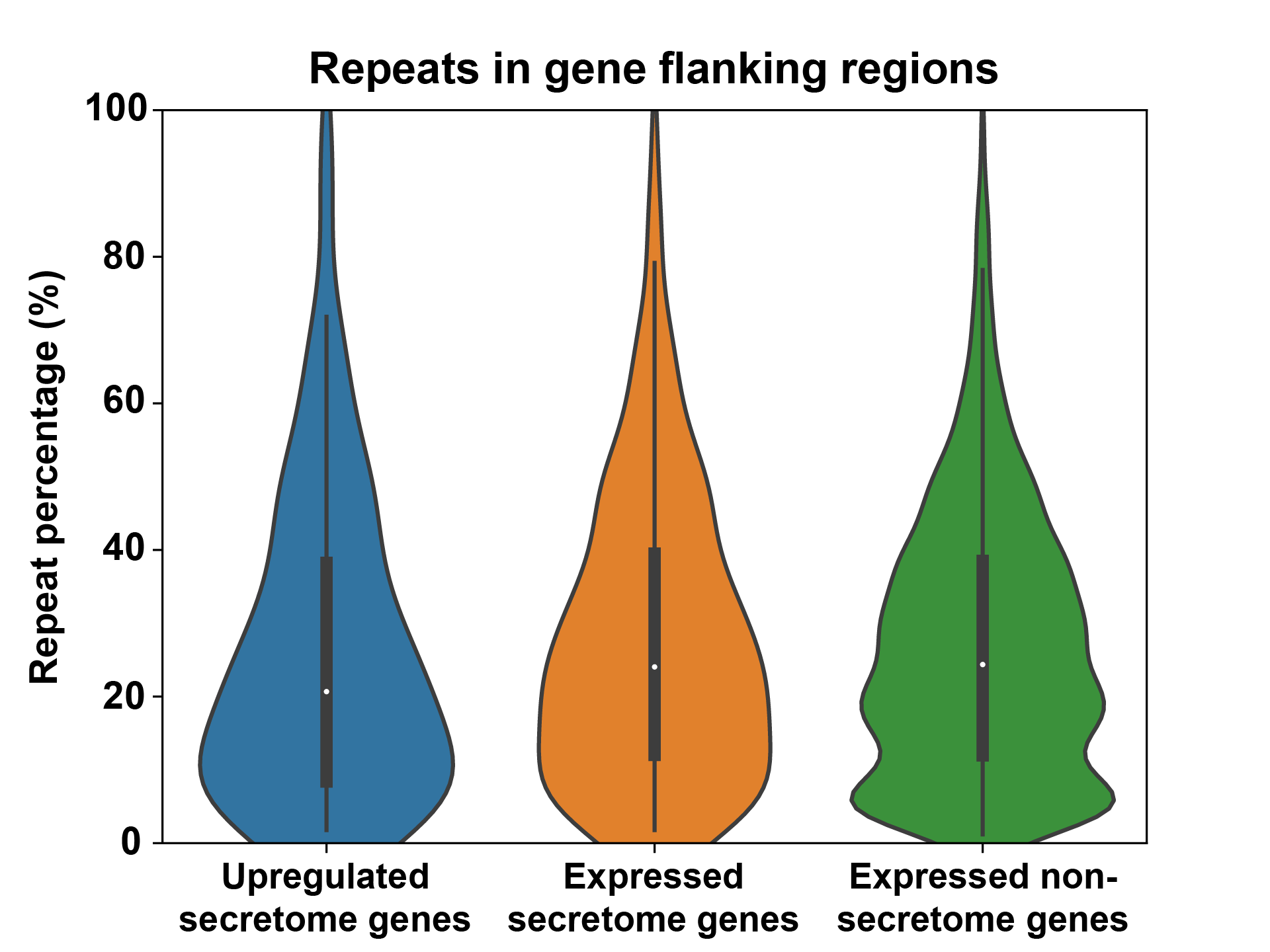

**Figure S2 Comparison of repeat content of gene flanking regions between expressed secretome genes and other expressed genes.** Upregulated means a significantly higher expression *in planta* compared to growth in axenic culture. In total, 20 kb sequences on each side of the genes (40 kb in total) were considered as flanking regions. Significant differences were calculated with a two-sided T-test. No significant differences with *p*-value < 0.01 were found.
