## Supplementary material for "High nucleotide substitution rates associated with retrotransposon proliferation drive dynamic secretome evolution in smut pathogens": Figure S3

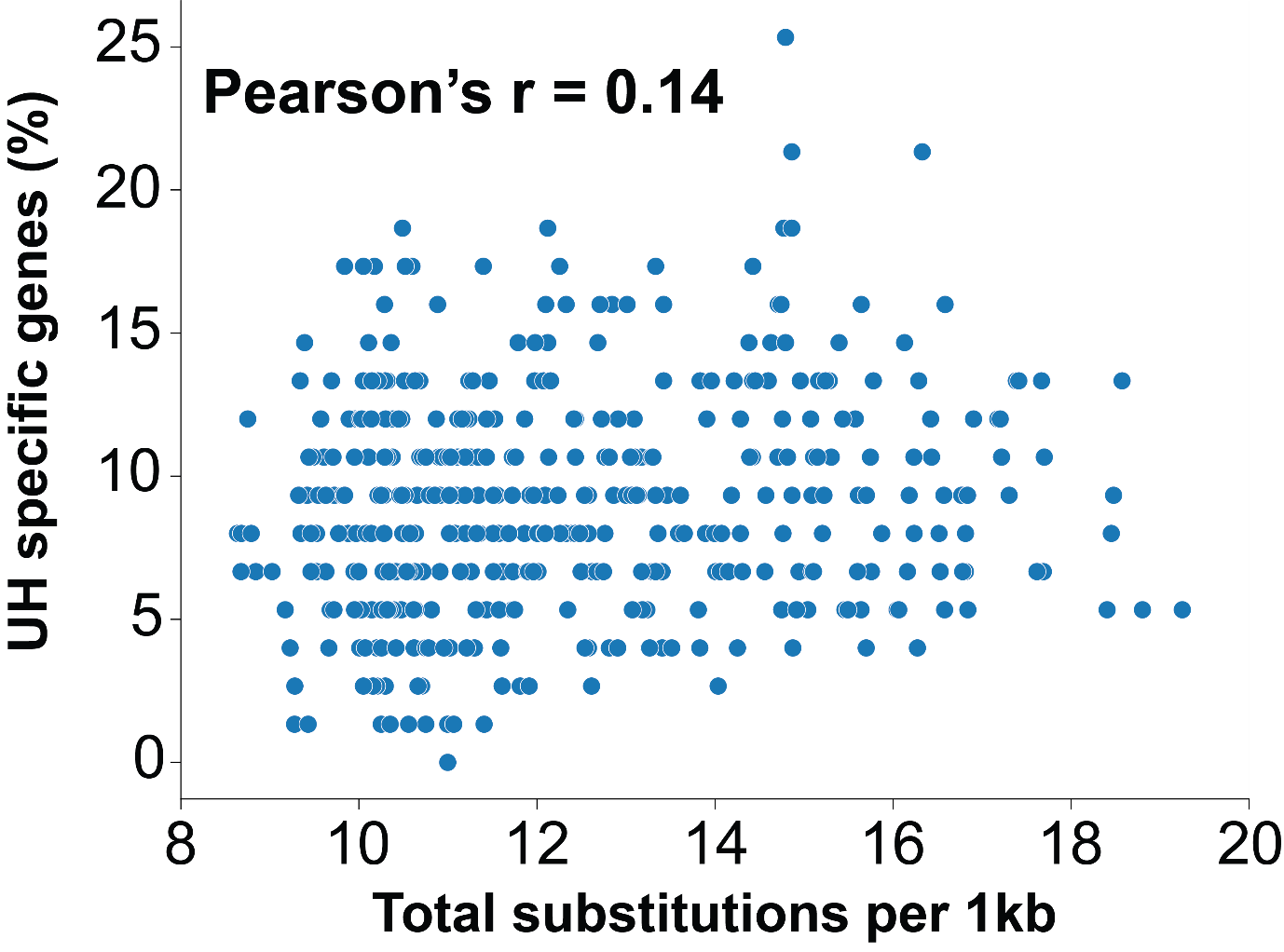

**Figure S3: Correlation between median nucleotide substitution level and fraction *U. hordei* (UH) specific genes for ortholog windows.** Ortholog windows of 75 UH genes with a sliding step of 10 were used to determine the number of substitutions with *U. nuda*. UH specific genes do not have an ortholog in *U. maydis*.
