## Supplementary material for "High nucleotide substitution rates associated with retrotransposon proliferation drive dynamic secretome evolution in smut pathogens": Figure S4

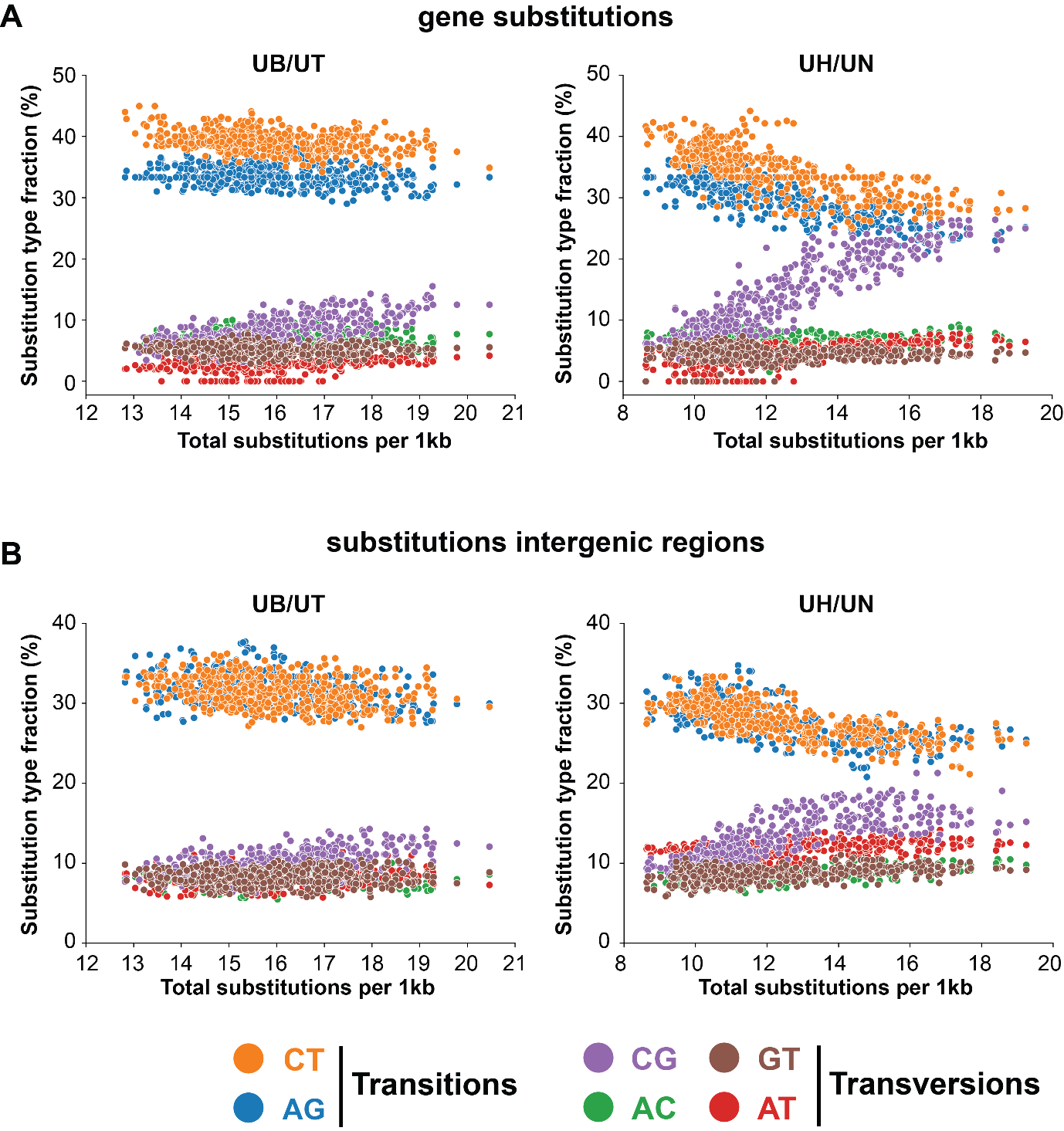

**Figure S4: Comparison of nucleotide substitution type fractions for *U.* *brachipodii-distachyi*/*U. tritici* and *U. hordei*/*U. nuda* ortholog windows.** The nucleotide substitutions were calculated for windows of 75 genes with a sliding step of 10. The x-axis consistently displays the total substitutions per 1 kb for these windows. (**A**) The y-axis depicts the fraction of every substation type (CT, AG, CG, AC, GT, AT) of ortholog windows. (**B**) The y-axis depicts the fraction of every substation type for the intergenetic regions of ortholog windows.
