## Supplementary material for "High nucleotide substitution rates associated with retrotransposon proliferation drive dynamic secretome evolution in smut pathogens": Figure S5

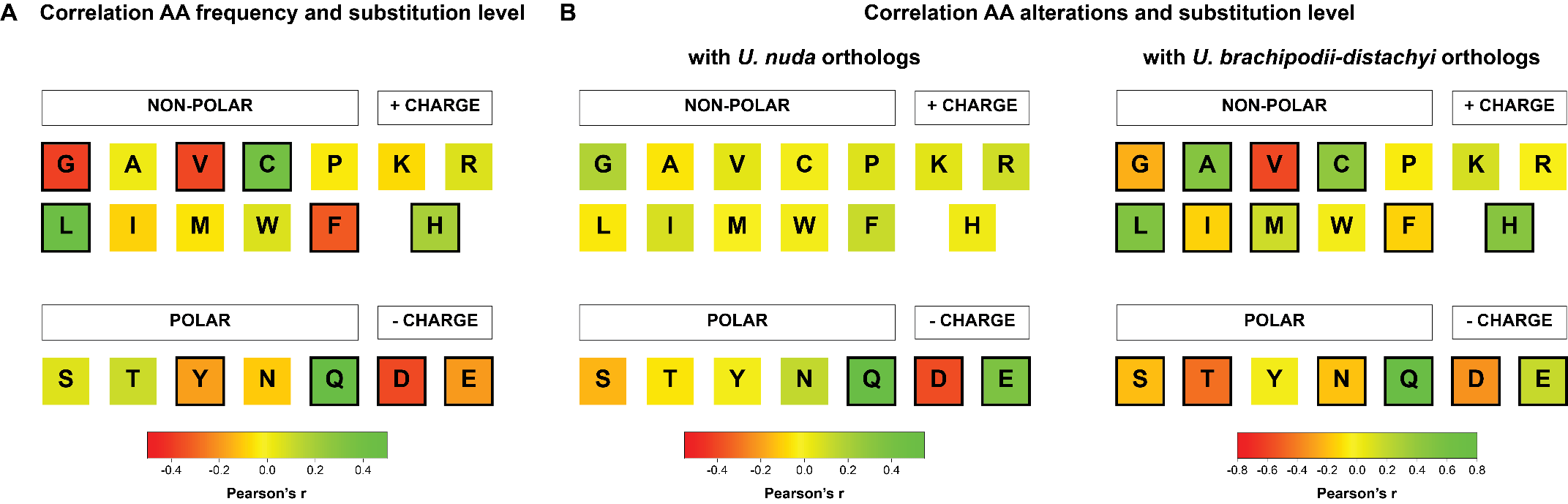

**Figure S5: Correlations between the nucleotide substitution levels and the encoded amino acid composition of genes.** Correlations were calculated for windows of 75 *U. hordei* genes with a sliding step of 10. Significant correlations, with *p*-value < 0.01, are indicated by a black edge around the square. (**A**) Correlations between amino acid compositions of encoded *U. hordei* proteins and the number of nucleotide substitutions with *U. nuda*. (**B**) Correlations between amino acid alternations for encoded *U. hordei* proteins and the number of nucleotide substitutions using *U. nuda* and *U. brachipodii-distachyi* orthologs as comparison.
