## Supplementary material for "High nucleotide substitution rates associated with retrotransposon proliferation drive dynamic secretome evolution in smut pathogens": Figure S6

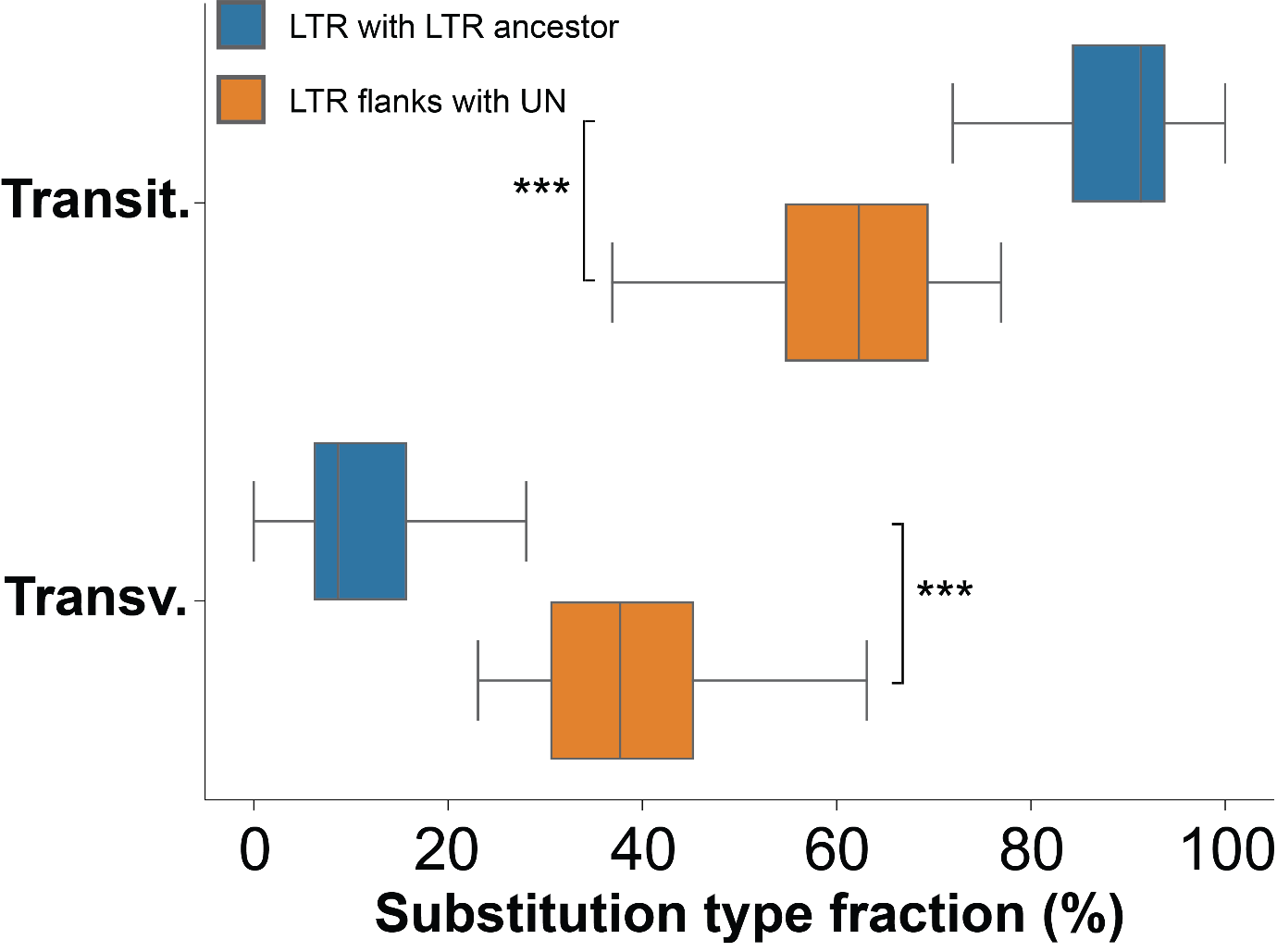

**Figure S6: Fractions of transitions and transversion of recently proliferated *U. hordei* long terminal repeat retrotransposons (LTR-RTs) and their flanking regions.** The fraction of transitions and transversions between recently proliferated *U. hordei* LTR-RTs and their ancestors were determined. Fractions of the 20 kb flanking regions (40 kb in total) of the LTR-RT with *U. nuda* (UN) were also determined. Significant differences between LTR and flanking regions were determined for transitions and transversions separately with an unequal variance t-test. ***: *p*-value < 0.001.
